# Reverse genetics in humanized mice reveals CARD8-mediated pyroptosis causing pancytopenia in human DPP9 deficiency

**DOI:** 10.64898/2026.06.27.735024

**Authors:** Tianli Xiao, J. Richard Brewer, Maximilian Carlino, Ailin Han, Yamato Takabe, Chia-Yi Lee, Fengrui Zhang, Mi Chen, Holly Nicole Blackburn, Amin H. Nassar, Qiankun Wang, Kristen Brennad, Liang Shan, Esen Sefik, Diane S. Krause, Richard A Flavell

## Abstract

Loss of function mutation in the human *DPP9* gene causes Hatipoglu syndrome leading to severe inflammasomopathy. A key feature of the disease is pancytopenia and patients require bone marrow transplantation, but the mechanism of cell loss is unclear since *Dpp9* mutant mice have normal hematopoiesis, suggesting that a distinct mechanism of disease occurs in humans. Here, we present a model of human DPP9 deficiency leveraging reverse genetics in the MISTRG6 humanized mice. We found that CRISPR editing of human CD34^+^ hematopoietic stem and progenitor cells (HSPCs) led to very efficient and persistent gene deletion in vivo. Human *DPP9* deletion recapitulated cytopenia in peripheral blood and in the bone marrow, and cell loss was cell intrinsic. However, *DPP9* deletion led to little transcriptional changes suggesting post-transcriptional regulation in human HSPCs. Mechanistically, DPP9 deficiency led to the activation of the CARD8 inflammasome resulting in HSPC pyroptosis, whereas NLRP1 was dispensable for cell death. Thus, our results reveal a unique human mechanism of disease and offer therapeutic insight for this inflammasomopathy.

## Introduction

Rare autoinflammatory syndromes arising from dysregulation of the inflammasome, or inflammasomopathies, have revealed important mechanisms of human innate immunity. Although many of the principles in inflammasome biology are shared between mouse and human, over 75 million years of distinct microbial exposure have imposed different evolutionary pressure on inflammasome sensors, raising the question whether some inflammasomopathies have distinct mechanisms between humans and model organisms such as mice.

Recently, an inflammasomopathy caused by loss-of-function mutations in the *DPP9* locus was identified in several families (1, 2). These patients present with recurrent fevers, repeated infections, pancytopenia and anemia, and elevated serum levels of inflammatory cytokines reported in one patient. Current treatment requires bone marrow transplantations (1). How human *DPP9* mutations cause pancytopenia remains an important question, since pancytopenia likely causes recurrent infection due to the lack of immune cells and the need for bone marrow transplant. Strikingly, this hematopoietic loss is not recapitulated in mice —*Dpp9* mutants in conventional mouse models maintain normal immune cell numbers, and their hematopoietic stem cells retain full reconstitution capacity upon serial transplantation (1, 3, 4). Rather than dismissing this discrepancy as a limitation of animal modeling, we reasoned that the divergent evolutionary trajectories of mouse and human inflammasome biology may have endowed human hematopoietic stem cells with a distinct, species-specific vulnerability, one that, if identified, could illuminate not only the pathogenesis of DPP9 deficiency, but a previously unrecognized axis of human blood stem cell regulation.

*DPP9* encodes dipeptidyl peptidase 9, which typically cleaves X-P dipeptides where X is any amino acid. It also serves as the endogenous inhibitor of mouse NLRP1, human NLRP1 and human CARD8 inflammasomes (5–10). NLRP1 and CARD8 are two structurally related inflammasomes that cause pyroptotic cell death and release IL-1β and IL-18 proinflammatory cytokines upon activation (7, 8, 11–14). Both NLRP1 and CARD8 contain a regulatory N-terminus and cytotoxic C-terminus. The cytotoxic C-terminal peptide contains a CARD domain that can interact with ASC or caspase-1 (CASP1) for inflammasome activation. However, spurious activation by these C-terminal peptides is prevented by DPP9 through the formation of a ternary complex (5, 6). During inflammasome activation, the N-terminal peptide undergoes “functional degradation” by the proteasome, which releases additional cytotoxic C-terminal fragments and overwhelms DPP9 complexes leading to inflammasome activation (15, 16). Importantly, NLRP1 and CARD8 are poorly conserved between mice and humans to the point that human and mouse NLRP1 are activated by distinct signals (17, 18). On the other hand, the CARD8 inflammasome is absent in most rodent species including *Mus musculus*, but is widely expressed in human hematopoietic cells(18, 19).

We address how human DPP9 deficiency causes pancytopenia in vivo by leveraging the MISTRG6 humanized mouse model. MISTRG6 mice enable the engraftment of human CD34^+^ hematopoietic stem and progenitor cells (HSPCs) by modification of several key genes to express human **M**-CSF(20, 21), **I**L-3 and GM-CSF(22), **S**IRPα(23), **T**hrombopoietin(24), IL-**6** (21, 25) in a ***R****ag**^-/-^***and *IL2r**g**^-/-^* background(26, 27). Physiologic expression of these human factors is achieved by gene knock-in replacement of their musine counterparts. The immunodeficient background, combined with expression of human factors, enable human CD34^+^ HSPCs to engraft and differentiate into a near complete complement of human immune system, including human NK cells, monocytes and macrophages, T cells and B cells. Human hematopoietic stem cells (HSC) persist long-term in the bone marrow of MISTRG6 mice and can be serially transplanted up to four generations (26–28). MISTRG6 and related family of strains have revealed mechanisms of human immunity in the context of infection, cancer, autoimmunity, and hematopoiesis (26, 28–39).

To resolve this species-specific paradox, we combined xenotransplantation, CRISPR gene editing, flow cytometry, colony forming unit assays, competitive transfers and single-cell transcriptomics, we found that human *DPP9^-/-^* HSPCs undergo pyroptosis through activation of the CARD8 inflammasome, which triggers caspase-1-dependent cell death. Deletion of *CARD8* or *CASP1* rescued cytopenia and loss of DPP9-deficient bone marrow stem cells, whereas NLRP1 deficiency did not. Given that *CARD8* is absent from the mouse genome yet highly enriched in human hematopoietic populations, we propose that the mouse-human discrepancy in DPP9 deficiency reflects a fundamentally distinct inflammasome logic governing human blood stem cell survival, one in which CARD8 serves as a sentinel whose unchecked activation is sufficient to collapse human hematopoiesis.

## Results

### Efficient and persistent gene knockout of human CD34+ HSPCs in the MISTRG6 humanized mouse

Since DPP9 deficiency has different phenotypes in mouse and human, we set out to establish a model of human DPP9 deficiency using the MISTRG6 humanized mouse system. To generate DPP9 deficiency or gene knock out in general, we initially tested the feasibility of generating CD34^+^ hematopoietic stem and progenitor cells (HSPC) from iPSCs using a protocol from Stemcell Technologies. However, iPSC-derived CD34^+^ failed to generate substantial (>1%) engraftment in MISTRG6 mice, despite expressing markers of human HSCs such as CD34, CD90 and CD49f. (Supp Fig. 1A-B). Therefore, we explored gene editing in primary human CD34^+^ HSPCs isolated from cord blood or fetal liver. We optimized the ratio and quantity of Cas9 and sgRNA to delete *TRAC* encoding the T cell receptor alpha chain (TCRα), which is required for T cell development, in human CD34^+^ HSPCs. Inference of CRISPR Edits analysis (40) revealed that CRISPR RNPs generated large truncations in the *TRAC* locus (Supp Fig. 1C-E). We further optimized culture conditions to promote CD34^+^ HSPC recovery post gene editing (Supp Fig. 1F). Next, we assessed whether edited cells could engraft in the MISTRG6 animals and whether gene knockouts persist in vivo. Immediately after electroporation, we administered edited CD34^+^ HSPCs or control cells, which were edited with control sgRNA RNPs, via intrahepatic injection to neonatal MISTRG6 pups (Figure 1A) without preconditioning. Deletion of *TRAC* was persistent in vivo, as mice engrafted with *TRAC^-/-^*human HSPCs failed to generate human T cells in the spleen, liver, and lung up to 16 weeks post engraftment, whereas control mice had abundant human T cells (Figure 1B and C). Development of other human hematopoietic cells were not impaired, preserving B cells and myeloid cells (Supp Fig. 1G-I). As an additional proof of concept, we tested deletion of human *CSF1R*. As expected, CSF1R expression was significantly decreased in human blood monocytes in MISTRG6 mice 9 weeks after engraftment (Figure 1D). Furthermore, the number of CD16^+^ monocytes and liver macrophages were significantly decreased in the blood (Figure 1E and F). MISTRG6 mice with *CSF1R^-/-^*HSPCs retained lymphoid cell development, and human CD45^+^ were similar in *CSF1R^-/-^* HSPCs (Supp Fig. 1J and K). Taken together, CRISPR RNPs led to highly efficient knockout in human CD34^+^ HSPCs that persist in vivo in MISTRG6 humanized mice.

**Figure 1.**
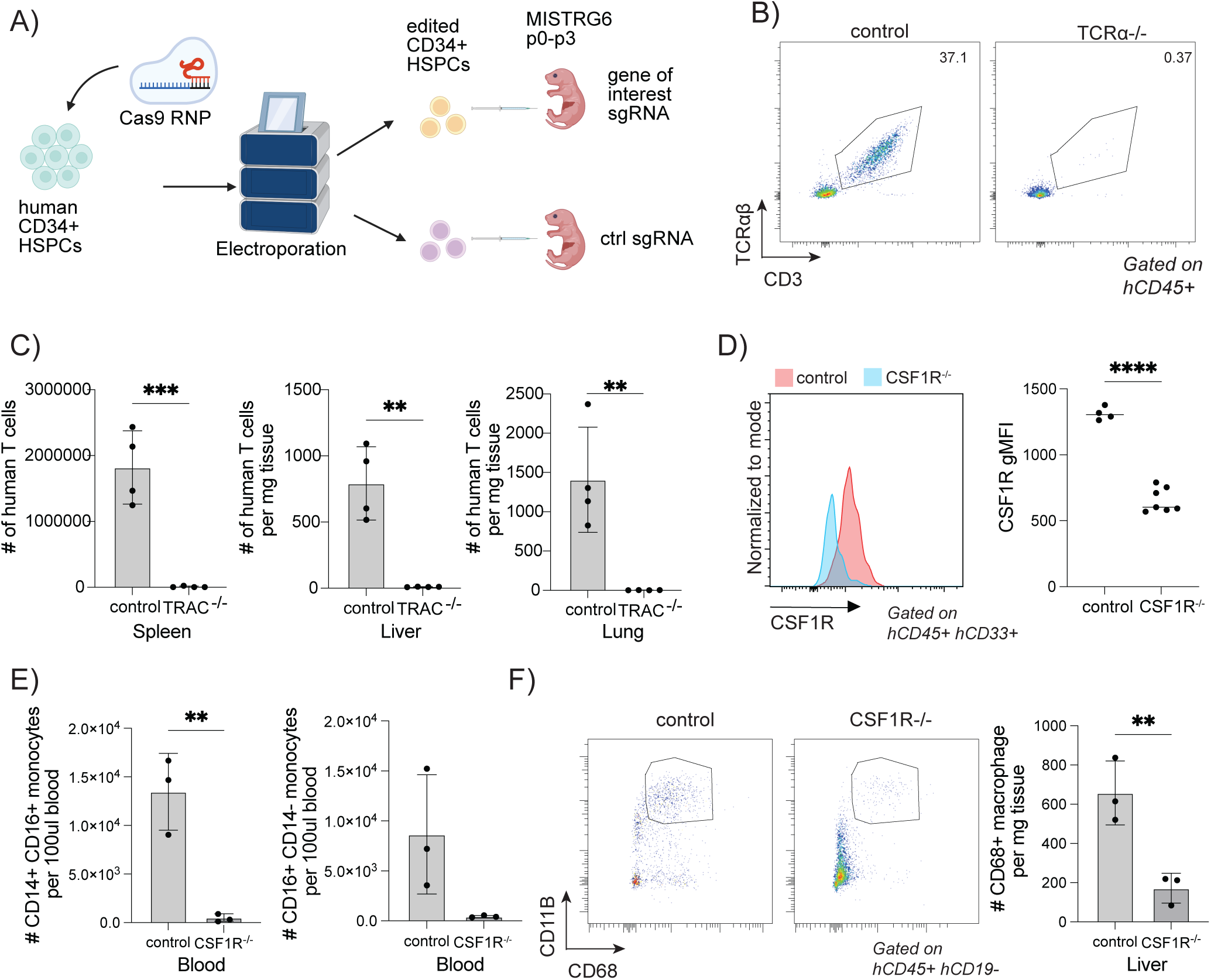
Efficient and persistent gene knockout of human CD34+ HSPCs in MISTRG6 humanized mouse model. (**A**) Schematic of gene editing followed by engraftment in the MISTRG6 humanized mouse model. MISTRG6 pups were injected intrahepatically in the first three days after birth without preconditioning. (**B-C**) *TRAC^-/-^*and control CD34^+^ HSPCs were engrafted in MISTRG6 mice and assessed 12 to 16 weeks after engraftment. Data pooled from two experiment. (**B**) Representative flow plot from spleen. (**C**) Number of human T cells in tissues. (**D-F**) *CSF1R^-/-^* and control CD34^+^ HSPCs were engrafted in MISTRG6 mice and assessed 9 weeks after engraftment. (**D**) Expression of CSF1R on human CD33+ myeloid cells in the blood 9 weeks after engraftment. Left: representative histogram. Right: summary data. (**E**) Number of cells in blood. (**F**) Number of cells in the liver. Experiments are representative from at least two human donors. *P ≤ 0.05, **P ≤ 0.01, ***P ≤ 0.001, and ****P ≤ 0.0001 according to Student’s t test.

### Human DPP9 deficiency leads to pancytopenia and bone marrow failure

To assess the consequence of human DPP9 deficiency in vivo, we designed three sgRNA targeting the catalytic exon of *DPP9* in CD34^+^ HSPCs and engrafted them via intrahepatic injection to MISTRG6 mice, then assessed their engraftment 8-9 weeks later. PCR amplification of the knockout region revealed efficient deletion in vitro and in vivo after engraftment (Supp Fig. 2A). Mice engrafted with *DPP9^-/-^* HSPCs (hereafter referred to as *DPP9^-/-^* mice) had significantly fewer human CD45^+^ cells in the blood, including monocytes and B cells (Figure 2A), thus recapitulating leukopenia in patients with *DPP9* germline mutations (1). T cell numbers were comparable between control and *DPP9^-/-^*mice, likely because once an initial wave of T cell progenitors differentiate into mature T cells and undergo homeostatic expansion. Once T cells become activated, they become resistant to CARD8-mediated cell death (41, 42). Furthermore, in many humanized mouse models, T cells preferentially expand in poorly engrafted mice and thus do not faithfully reflect stem cell differentiation in the bone marrow (43). The phenotype was consistent between human donors (Figure 2A). We reasoned that peripheral cytopenia may be due to a loss of stem cells in the bone marrow. Indeed, *DPP9^-/-^*Lin^-^ CD34^+^ HSPCs failed to persist in the bone marrow (Figure 2B). Further dissection of CD34^+^ population revealed that the number of common lymphoid progenitor (CLP), common myeloid progenitor (CMP), megakaryocyte-erythroid progenitor (MEP), hematopoietic stem cells (HSCs), and multipotent progenitors (MPP)s were significantly reduced in *DPP9^-/-^*mice (Supp Fig. 2B, Figure 2C). To ensure that loss of *DPP9^-/-^* HSPCs was not due to their inability to home to the bone marrow, we assessed whether direct intrafemoral injection of *DPP9^-/-^* HSPCs could ameliorate engraftment in vivo. Despite direct access to bone marrow space, *DPP9^-/-^*HSPCs failed to persist in the bone marrow, and cytopenia was still observed in the blood in those animals (Supp Fig. 2C).

**Figure 2.**
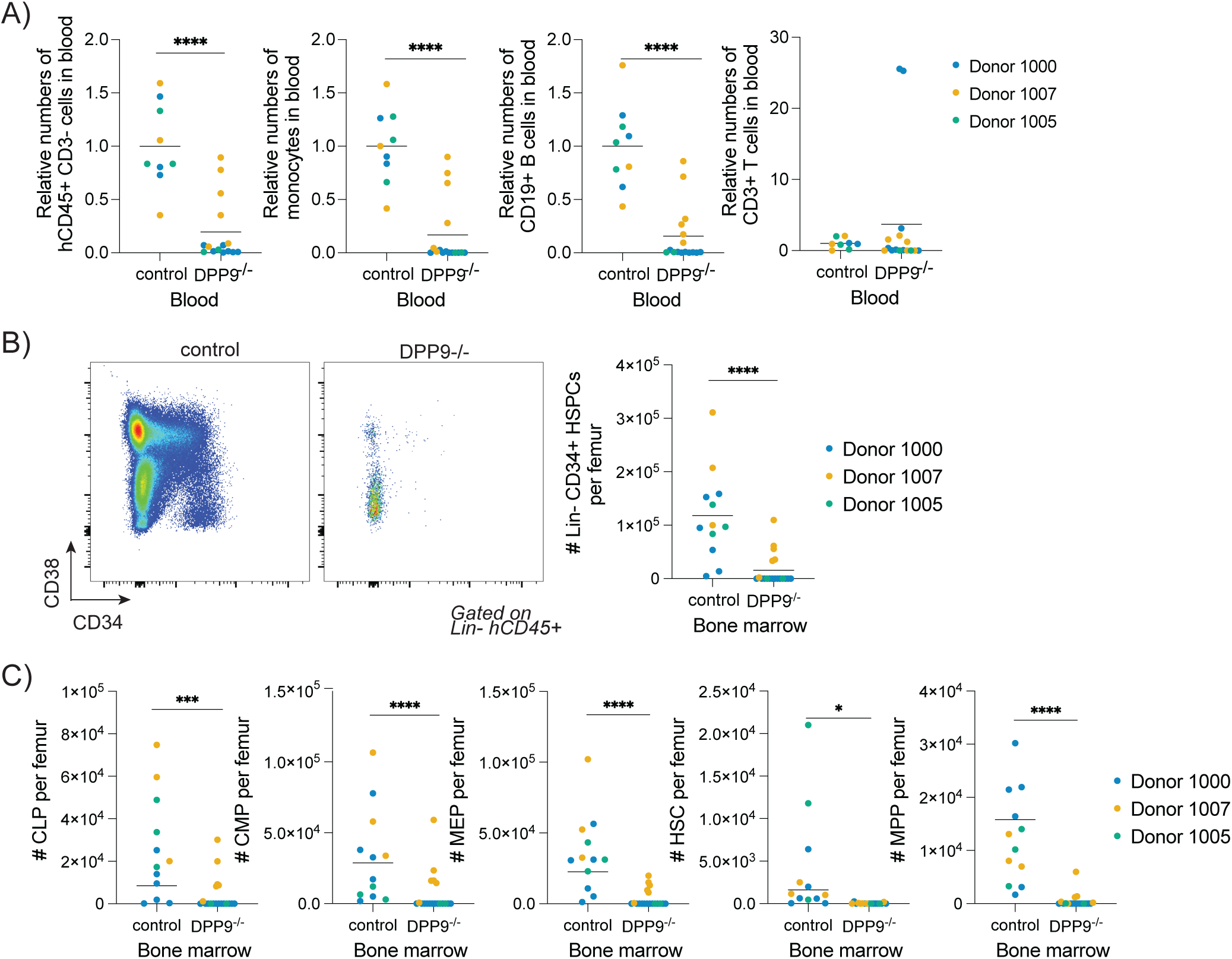
Human DPP9 deficiency results in cytopenia and loss of bone marrow hematopoietic stem and progenitor cells. MISTRG6 mice are engrafted with control or *DPP9^-/-^*CD34^+^ HSPCs for 8 to 9 weeks. (**A**) Relative number of cells in the blood. (**B**) Representative flow plot (left) and summary data (right) of lineage^-^ CD34^+^ HSPCs in the bone marrow. (**C**) Number of CD34^+^ HSPC subsets in the bone marrow. CLP, common lymphoid progenitor. CMP, common myeloid progenitor. MEP, megakaryocyte-erythroid progenitor. HSC, hematopoietic stem cell. MPP, multipotent progenitor. Relative cell numbers are calculated by normalizing cell counts of each animal to the mean value of the control group per donor and per engraftment in MISTRG6 mice. *P ≤ 0.05, **P ≤ 0.01, ***P ≤ 0.001, and ****P ≤ 0.0001 according to 2-way ANOVA between genotypes.

### Loss of CD34^+^ *DPP9^-/-^* HSPCs is cell-intrinsic

To assess whether loss of *DPP9^-/-^* HSPCs is recapitulated in vitro, we knocked out DPP9 and cultured control (*TRAC^-/-^*) and *DPP9^-/-^* HSPCs in liquid culture. Surprisingly, *DPP9^-/-^*HSPCs expanded normally in vitro (Supp Fig. 3A). We reasoned that defects with DPP9 mutants would be more prominent at a clonal level, thus we seeded control and *DPP9^-/-^*CD34^+^ cells in a colony forming unit (CFU) assay utilizing Megacult medium supplemented with cytokines supportive of granulocyte, monocyte, megakaryocyte, and erythroid lineages to detect any specific lineage biases (Figure 3A-C). We observed fewer colonies formed by *DPP9^-/-^* HSPCs (Figure 3B) although the effect was much more modest than loss of HSPCs in vivo. DPP9 mutation did not lead to lineage bias (Figure 3C), consistent with our findings in vivo that all lineages were affected by DPP9 mutation (Figure 2C). To specifically assess the impact of DPP9 deficiency on hematopoietic stem cells (HSC), we sorted single CD34^+^ CD38^-^ CD90^+^ human HSCs into a 384-well plate and cultured them with either an expansion medium or differentiation medium (Figure 3D). Similar to bulk CD34^+^ populations, *DPP9^-/-^*HSCs expanded to fewer cells at the end of 7-day expansion (Figure 3E), and the percentage of clones that had substantial expansion (>100 cells) was also lower in *DPP9^-/-^* HSCs (Supp Fig. 3B). In contrast, differentiation into myeloid and erythroid lineages were similar between control and *DPP9^-/-^* HSCs (Figure 3F). Finally, to determine whether loss of *DPP9^-/-^* HSPCs in vivo is cell intrinsic, we engrafted a 1:1 mixture of control and *DPP9^-/-^* HSPCs in MISTRG6 mice and assessed the frequency of *DPP9^-/-^* HSPCs after 7 weeks (Figure 3G). Unlike competitive transfer assays in conventional mouse models where each donor is congenically marked, control and *DPP9^-/-^* HSPCs from the same human donor would not express different cell-surface markers. To distinguish the cells derived from the two genotypes, we leveraged digital droplet PCR to detect the fraction of control and knockout cells (39, 44–48). Thus, we sorted various bone marrow population from the competitive transfer recipient mice and found that *DPP9^-/-^*cells are nearly completely lost in the HSCs, progenitor cells and differentiated lin^+^ cells (Figure 3H). These data suggest that loss of *DPP9^-/-^* occurs in a cell-intrinsic manner.

**Figure 3.**
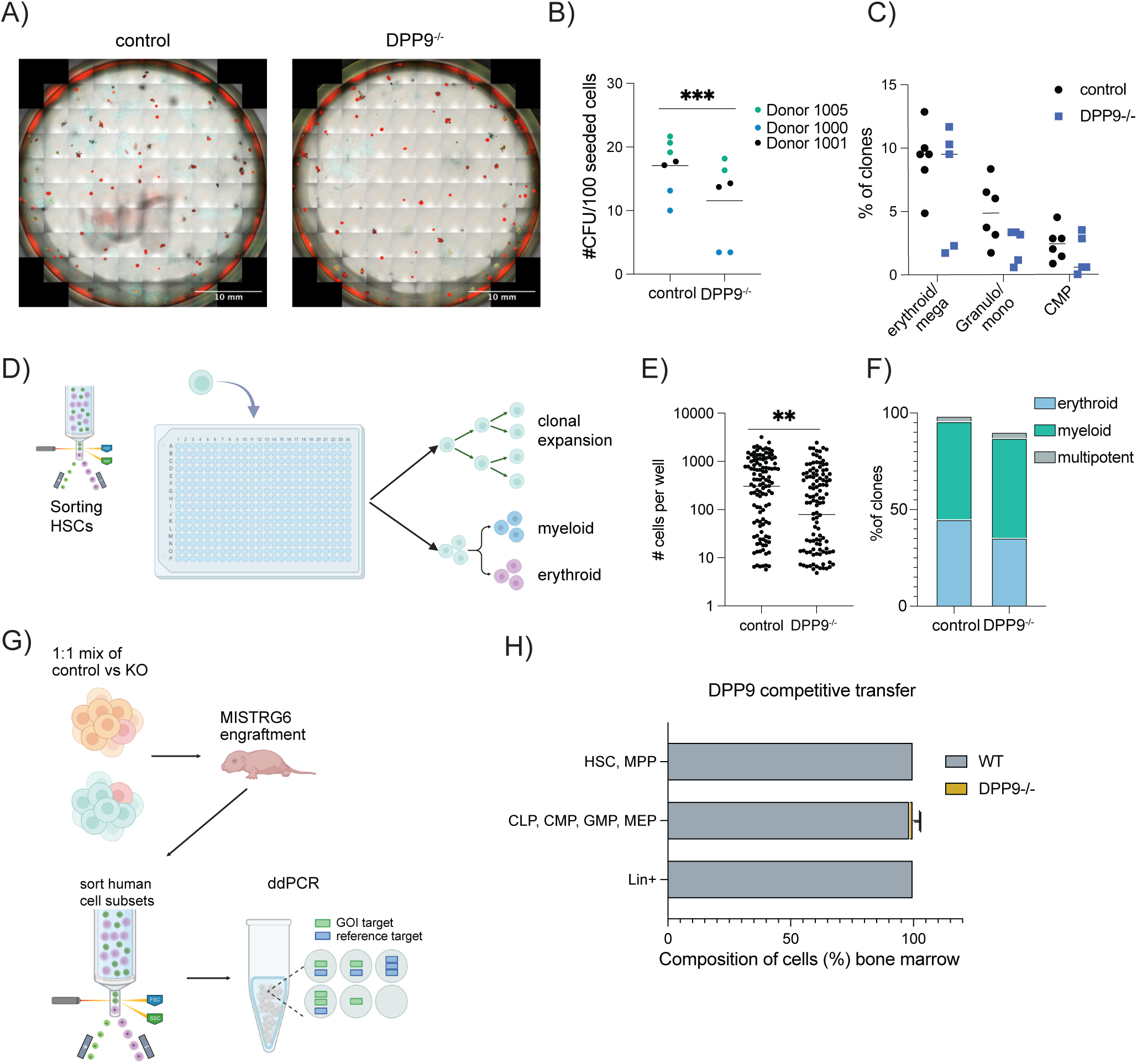
Loss of CD34+ *DPP9^-/-^*HSPCs is cell-intrinsic. (**A**) In vitro CFU assay plate. Red:CD235a; yellow: CD66b; cyan: CD14; green: CD41. (**B**) Summary of in vitro CFU assay. (**C**) Lineage differentiation from in vitro CFU assay. (**D**) Schematic of sorting single hCD45^+^ CD34^+^ Lin^-^ CD38^-^ CD90^+^ HSCs into 384-well plate for in vitro expansion and differentiation assay. (**E**) Number of cells per well after 7 day expansion of sorted HSCs. (**F**) Differentiation toward myeloid (CD66b^+^ or CD14^+^) and erythroid (CD235a^+^) lineages from sorted HSCs. Clones that give rise to both fates are labeled multipotent. (**G**) schematic of competitive transfer experiment in MISTRG6 mice. (**H**) Composition of human bone marrow cells 7 weeks after competitive transfer. CFU, colony forming unit. ddPCR, digital droplet PCR. CLP, common lymphoid progenitor. CMP, common myeloid progenitor. GMP, granulo-macrophage progenitor. MEP, megakaryocyte-erythroid progenitor. HSC, hematopoietic stem cell. MPP, multipotent progenitor. *P ≤ 0.05, **P ≤ 0.01, ***P ≤ 0.001, and ****P ≤ 0.0001 according to Student’s t test.

### DPP9 deficiency leads to few transcriptional changes in HSPCs

As a preliminary step to mechanistically understand how *DPP9^-/-^*HSPCs are lost in vivo, we performed single-cell (sc)RNA-seq. We generated both control and *DPP9^-/-^* HSPCs from two human donors and engrafted littermate MISTRG6 animals as above. Four weeks after engraftment at a time point when some *DPP9^-/-^* cells are still present, we sorted Lin^-^ CD34^+^ cells. Control and *DPP9^-/-^*HSPCs were separately hashtagged and encapsulated together. After demultiplexing, we obtained robust scRNA-seq data from over 5000 cells with an average of 5000 genes per cell. Dimensional reduction using Uniform Manifold Approximation and Projection (UMAP) and cluster analysis revealed 12 distinct clusters (Figure 4A). To identify them, we iteratively overlaid the signature from a large human bone marrow dataset (49) onto our clusters and identified two HSC/MPP clusters, three myeloid progenitor clusters, two common lymphoid progenitor clusters, five lymphoid progenitor clusters, and one erythroid progenitor cluster (Figure 4A, Supp Fig. 4A). We confirmed that each cluster expressed specific markers of their population (Figure 4B, Supplemental Table 1). For example, the HSC/MPP clusters highly expressed *CD34, AVP, ITGA6* (encoding CD49f) which are canonical markers of human HSCs (Supp Fig. 4B). Similarly, common lymphoid progenitor expressed *CD2*, whereas the pro-B cell progenitors expressed *CD81* and *CD72.* On the other hand, myeloid progenitors expressed *PRTN3, AZU1, MPO, and ELANE.* Megakaryocyte and erythroid progenitors expressed *SLC40A1* and *APOC1* (Figure 4B). Notably, the frequency of *DPP9^-/-^* cells was relatively decreased in the HSC/MPP clusters consistent with their loss (Supp Fig. 4C).

**Figure 4.**
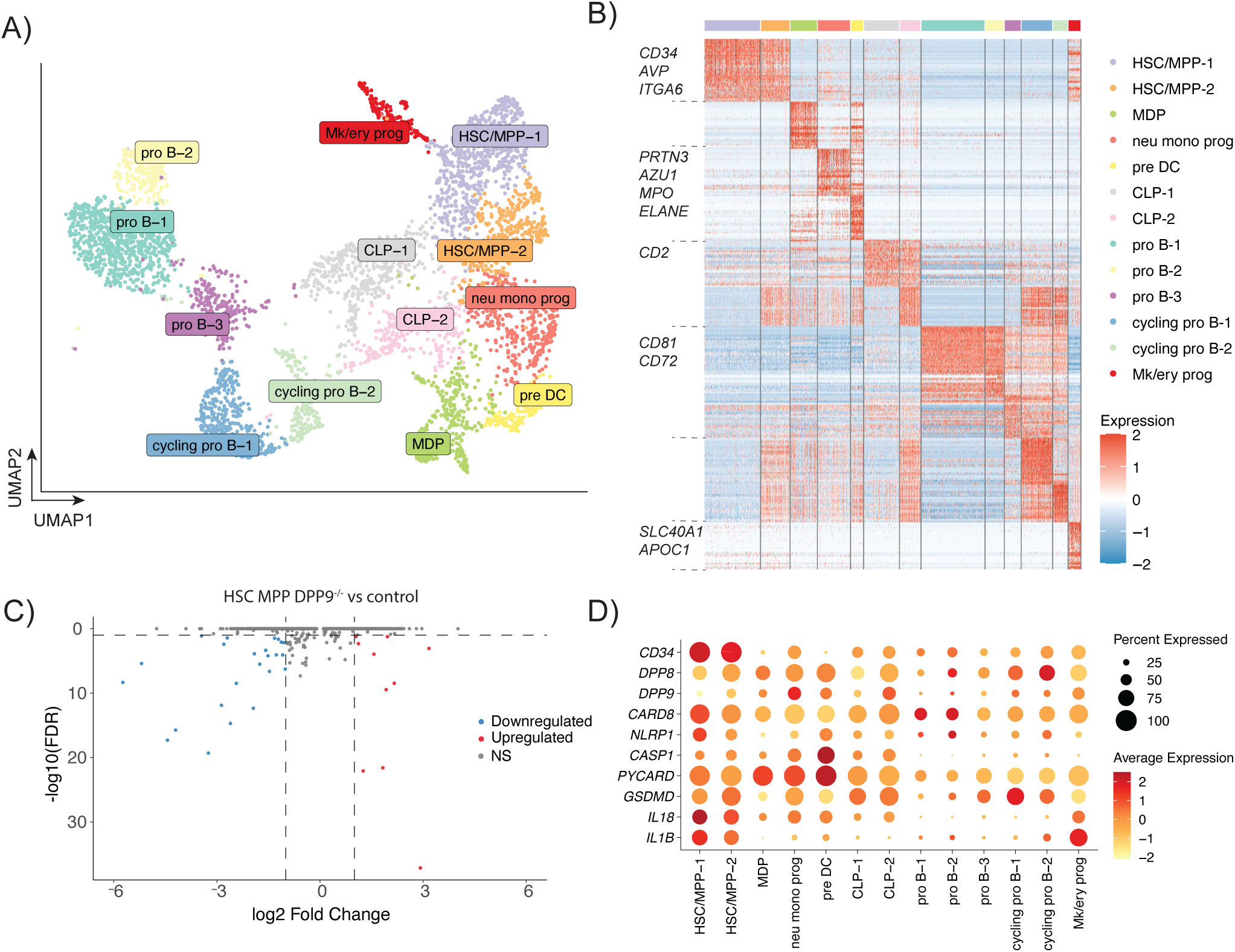
DPP9 deficiency in human CD34^+^ HSPCs does not result in drastic transcriptional changes. scRNA-seq analysis of sorted human Lin^-^ CD34^+^ cells from the bone marrow of MISTRG6 mice engrafted with either control or *DPP9^-/-^*CD34^+^ HSPCs, 4 weeks after engraftment. (**A**) Heterogeneity of human Lin^-^ CD34^+^ visualized by UMAP. (**B**) Heatmap of diagnostic genes of each cluster from the UMAP in (A). Top 30 genes are plotted for ease of visualization. (**C**) Volcano plot of differentially expressed (DE) genes between *DPP9^-/-^* and control cells from HSC/MPP-1 and HSC/MPP-2 clusters. (**D**) Gene expression of various inflammasome components across various human stem and progenitor subsets. CD34 expression (top) serves as a reference of a highly expressed gene.

To assess transcriptional changes in *DPP9^-/-^* cells, we performed differential expression analysis between control and *DPP9^-/-^*cells. We combined clusters of similar identity together to reveal potential subtle differences between genotypes, and combined the two HSC/MPP clusters (HSC/MPP-1, HSC/MPP-2), the three myeloid progenitor clusters (MDP, neu mono prog, pre DC), the two CLP clusters (CLP-1, CLP-2), and pro B cell clusters (pro B-1, pro B-2, pro B-3). Since DPP9 deficiency led to a loss of HSCs in vivo (Figure 2C), we focused our analysis on the HSC/MPP clusters. We detected only 77 differentially expressed (DE) genes in HSC and MPP cells (Figure 4C, Supp Fig. 4C, Supplemental Table 1). Ingenuity Pathway Analysis revealed that several signaling pathways were downregulated in *DPP9^-/-^* HSC and MPPs, which were primarily due to downregulation of *FOS* and *JUN*. Similarly, we found only 29 DE genes in myeloid progenitor cells, 14 DE genes in CLPs, and 16 DE genes in pro B cell clusters (Supp Fig. 4E, Supplemental Table 1). Taken together, scRNA-seq revealed that DPP9 deficiency did not lead to substantial transcriptional changes across any hematopoietic progenitor population, indicating that DPP9 controls HSPC fate primarily through post-transcriptional mechanisms. DPP9’s function as a protein-level inhibitor of the CARD8 and NLRP1 inflammasomes raises the possibility that unrestrained inflammasome activity, rather than any transcriptional reprogramming, underlies the progressive loss of DPP9-deficient HSPCs. To evaluate this, we first asked whether the relevant inflammasome machinery is even present in human HSPCs during xenotransplantation. Strikingly, *CARD8* was broadly and robustly expressed across CD34+ progenitor subsets, while *NLRP1* expression, though detectable, was considerably lower. Downstream effectors including *CASP1* and *GSDMD* were similarly expressed, indicating that human HSPCs possess the complete molecular apparatus required for inflammasome-driven pyroptosis (Figure 4D). *CARD8*’s absence in the mouse genome combined with the species-specific nature of the hematopoietic defect, pointed directly to CARD8 as the candidate mediator of DPP9-deficient HSPC loss and prompted us to test this genetically.

### CARD8 mediates pyroptosis of *DPP9^-/-^* HSPCs

To determine whether DPP9 deficiency leads to CARD8-mediated pyroptosis in CD34^+^ HSPCs, we cultured edited CD34^+^ HSPCs for three days and stimulated them with Val-boropro (VbP), an inhibitor of DPP8 and DPP9(7, 50). VbP treatment led to HSPC pyroptosis as measured by lactate dehydrogenase release. Notably, VbP elicited even higher pyroptosis in *DPP9^-/-^* HSPCs suggesting that DPP8 also plays a role in sequestering the inflammasome from activation. Pyroptosis was completely abrogated in HSPCs that lack *CARD8* or *CASP1* suggesting that HSPCs undergo CARD8-dependent pyroptosis (Figure 5A). To determine whether CARD8 mediated loss of *DPP9^-/-^* HSPCs in vivo, we performed double knockout experiments to determine whether loss of CARD8 rescues *DPP9^-/-^*HSPCs. We found that simultaneous gene knockouts led to high knockout efficiency without marked loss of cell numbers (Supp Fig. 5B and C). We engrafted control, *DPP9^-/-^*, *DPP9^-/-^ CARD8^-/-^*and *DPP9^-/-^ NLRP1^-/-^* HSPCs from the same donor into littermate MISTRG6 mice. As expected, *DPP9^-/-^* HSPCs were lost after engraftment. HSPC numbers were rescued by deletion of *CARD8*, suggesting that CARD8 mediated pyroptosis of *DPP9^-/-^* HSPCs in vivo, including HSCs and MPPs (Figure 5B-C). Interestingly, *NLRP1* deletion failed to rescue *DPP9^-/-^* HSPCs. (Figure 5B-C). Leukopenia in the blood was also prevented, as the number of human CD45^+^ cells, including monocytes and B cells, recovered after deletion of *CARD8* but not *NLRP1* (Figure 5D). To examine whether *DPP9^-/-^* HSPCs underwent pyroptosis, we deleted *CASP1* which also rescued the loss of *DPP9^-/-^* HSPCs (Figure 5E). Taken together, these results reveal that *DPP9^-/-^* HSPCs undergo CARD8-mediated pyroptosis after transplantation, and loss of bone marrow stem cells lead to peripheral cytopenia.

**Figure 5.**
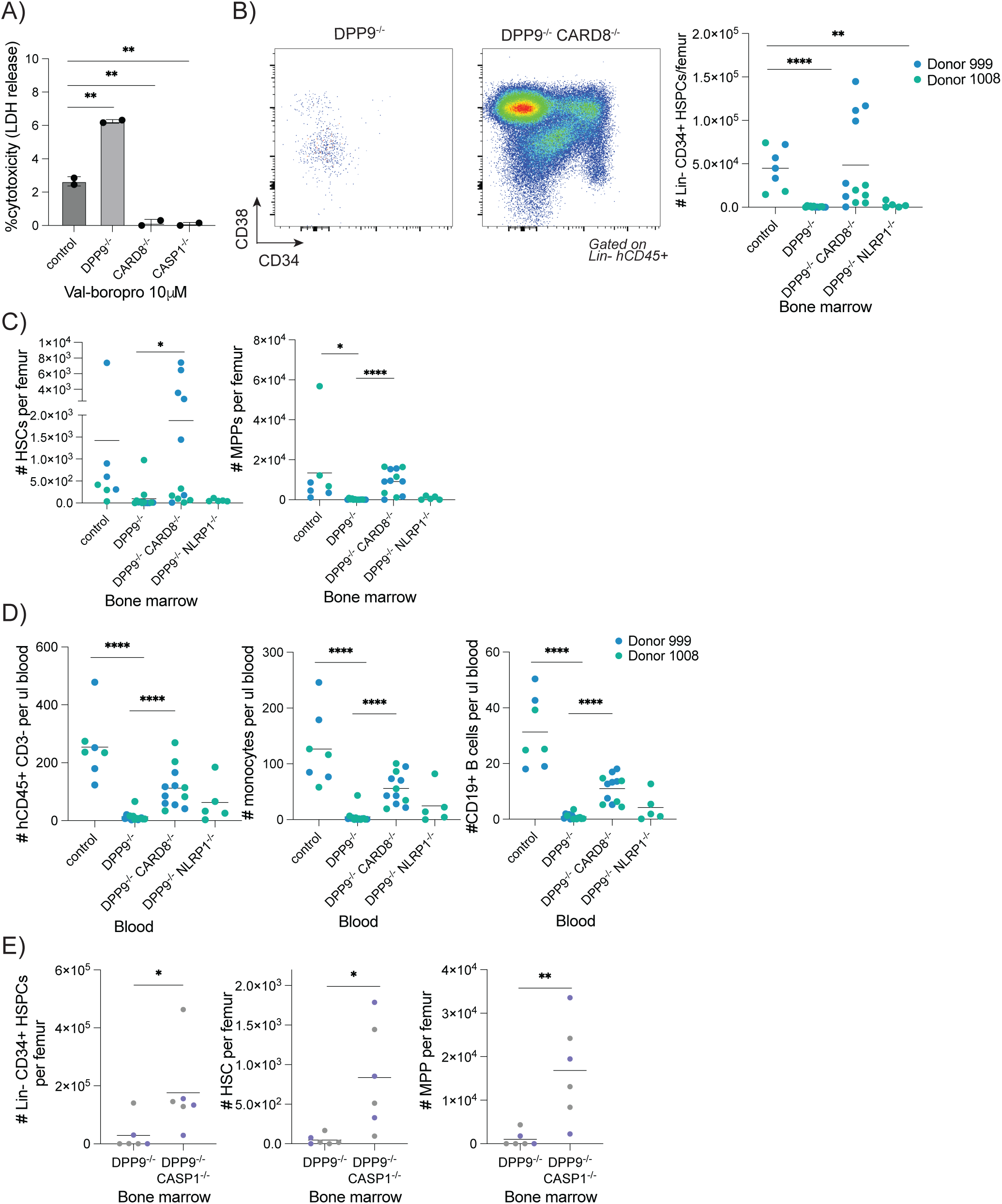
CARD8 mediates loss of *DPP9^-/-^* HSPCs. (**A**) Human CD34^+^ HSPCs are treated with Val-boroPro for 20 hours and lactate dehydrogenase was measured. (**B-D**) MISTRG6 mice were engrafted for 10 to 11 weeks. Cells of bone marrow (**B** and **C**) and blood (**D**) were quantified. (**E**) MISTRG6 mice were engrafted for 9 to 11 weeks. LDH, lactate dehydrogenase. *P ≤ 0.05, **P ≤ 0.01, ***P ≤ 0.001, and ****P ≤ 0.0001 according to Student’s t test.

## Discussion

The absence of a hematopoietic phenotype in conventional mouse models of *Dpp9* mutations contrasts with the devastating pancytopenia observed in patients. *Dpp9* mutant mouse HSCs are normal, since competitive transfer of wildtype and *Dpp9* mutant fetal liver cells leads to equal reconstitution in recipient mice and after secondary transplantation (3, 4). Additionally, the presence or absence of mouse NLRP1 inflammasome did not alter the number of bone marrow leukocytes in *Dpp9* mutant mice (1). Rather than reflecting a failure of the mouse model, we argue this discrepancy implies that the human hematopoietic compartment has acquired a distinct inflammasome-dependent vulnerability. *DPP9* mutations in patients lead to enzymatic hypomorphs, decreased protein expression or premature translation stop (1, 2), which we modeled in human HSPCs by generating a gene knockout targeting the *DPP9* catalytic exon. Our experiments reveal multi-lineage loss in the bone marrow and blood. The recapitulation of patient disease and the cell-intrinsic loss of stem cells suggest that the susceptibility to pyroptosis is a property of HSPCs, rather than a consequence of transplantation models. The near-absence of transcriptional changes in DPP9-deficient HSPCs further reinforces that DPP9 operates principally as a post-transcriptional gatekeeper. Our results indicate that DPP9 sets the activation threshold of the CARD8 inflammasome in human blood stem cells, and when this regulation is disrupted, triggers their depletion.

The hyperactivation of CARD8 inflammasome in human HSPCs in our in vivo model aligns with prior in vitro experiments which revealed increased NLRP1 and CARD8 activation in cells carrying *DPP9* mutations found in patients (1, 2). However, the selective requirement for CARD8, rather than NLRP1, in mediating the death of DPP9-deficient HSPC remains poorly understood. One possibility is that human DPP8 was sufficient to prevent NLRP1, but not CARD8, activation in *DPP9^-/-^* HSPCs. Indeed, in human keratinocytes, knockdown of both DPP8 and DPP9 is required for efficient NLRP1 activation (10). A second possibility is that *CARD8* is preferentially expressed in human HSPCs, unlike *NLRP1* which is highly expressed in barrier tissue cells in human (17, 51, 52). Indeed, we found that *CARD8* was well expressed in various stem and progenitor populations, whereas *NLRP1* is transcriptionally expressed at a lower level. Finally, a third possibility is that CARD8 is activated in the bone marrow microenvironment. Distinct and shared stressors activate human NLRP1 and CARD8 inflammasomes. Specifically, human NLRP1 is activated by ribotoxic stress (13, 14, 53, 54), viral RNA(55–57), toxins(58, 59), and viral proteases(15, 60–63). Human CARD8 is activated by various viral proteases and notably the HIV protease which induces pyroptosis in human T cells (11, 12, 19, 41). Recently, both NLRP1 and CARD8 were described to be activated by proteotoxic stress and reductive stress (50, 64–67). CARD8 may be triggered by a yet-to-be-defined cellular stress signal during transplantation of HSPCs and induce their pyroptosis in the absence of DPP9. How cell types determine whether to activate NLRP1 or CARD8 inflammasome will require further study. We also noted that the loss of *DPP9^-/-^* human HSPCs was far more pronounced in vivo than in vitro. This disparity may stem from differing experimental timeframes—over seven weeks in vivo versus less than two weeks in vitro—or it could suggest that in vivo signals actively accelerate *DPP9^-/-^* HSPC loss. For example, stem cells may compete for limiting quantities of survival and expansion signals such as human TPO within the bone marrow niche. They may also accumulate proliferation-induced protein folding stress capable of triggering CARD8 activation, as discussed above(50). Overall, the phenotypic differences in vivo and in vitro reinforces the value of using in vivo humanized mouse models to study human stem cell regulation.

Unlike human *NLRP1*, whose gain-of-function mutations cause skin pathology (68) reflective of its high expression in barrier cells, gain-of-function mutations of mouse *Nlrp1* causes pyroptosis of macrophage progenitor and granulocyte-macrophage progenitor cells. These mice have decreased numbers of lymphoid, myeloid, and erythroid cells in addition to systemic inflammation driven by IL-1β and IL-18 (69). Notably, mouse *Nlrp1* is highly expressed in hematopoietic lineages (69) akin to CARD8’s expression pattern in humans. Thus, human CARD8 and mouse NLRP1 may share some functional similarity, although each inflammasome maybe distinctly regulated owing to structure and sequence differences between these sensors (18).

Beyond DPP9 biology, our study establishes a generalizable platform for interrogating human-specific gene function in vivo. By generating control and knockout cells from the same human donor and using littermate MISTRG6 mice as recipients, we avoid confounding effects of diverse human genetics between donors and isolate the role of specific human genes in vivo (26, 70, 71). Thus, this approach can be applied to understand the role of other human genes that is poorly modeled in conventional mouse models. Given that SNP variants of *DPP9* were recently identified to regulate outcomes of idiopathic pulmonary fibrosis and SARSCoV2 (72–74), our study highlights the role of human DPP9 in restraining CARD8 inflammasome activation in human hematopoietic cells and contribute to our understanding of this emerging axis in human inflammatory diseases.

## Methods

### Sex as a biological variable

This study uses both male and female human CD34^+^ donor cells and both male and female recipient MISTRG6 mice. Similar findings are reported for both sexes.

### Mice and engraftment

MISTRG6 mice are previously described. Briefly, mice were generated by combining strains made by the R. Flavell lab, M. Manz lab and Regeneron Pharmaceuticals based on *Rag2*^−/−^ *IL2rg*^−/−^ 129xBalb/c background, followed by additional gene knock-in of human *CSF1*, *IL3/CSF2*, *SIRPA*, *THPO* and *IL6* in their respective mouse loci (21, 25, 26). Experimental mice are obtained by crossing MITRG6 with MISTRG6 to obtain MIS^h/m^TRG6, such that SIRPα is heterozygous for mouse and human, and M-CSF, IL-3/GMCSF, Thrombopoietin, IL-6 are homozygous for the human gene. For simplicity, these MIS^h/m^TRG6 recipients are referred to as MISTRG6 in all other sections of this manuscript. For engraftment, newborn MIS^h/m^TRG6 mice between 1 and 3 days old are injected intrahepatically with 30,000 purified CD34^+^ human hematopoietic stem and progenitor cells resuspended in 20ul using a 31G insulin syringe (BD). Neonatal MIS^h/m^TRG6 mice are not pre-conditioned prior to engraftment. Engrafted pups were cross fostered by CD1 dames until weaning. CD1 Elite mice are purchased from Charles River. Both male and female mice were used for engraftment. For intrafemoral engraftment, adult MIS^h/m^TRG6 mice were sublethally irradiated at 150 RAD (X-ray with X-RAD 320 irradiator). Mice were anesthetized under isoflurane, and a hole is made in the femur followed by injection containing 30,000 CD34^+^ HSPCs. Appropriate littermate controls were always used. All animals were housed in specific pathogen-free facilities at Yale Animal Resource Center and all procedures were carried out following protocols approved by the Institutional Animal Care and Use Committee.

### Isolation of CD34^+^ HSPCs

CD34^+^ cells are isolated from fetal liver purchased from Celce (previously known as Advance Bioscience Resources) or from cord blood collected by the Yale University Reproductive Sciences Biobank. CD34^+^ HSPC isolation from fetal liver: liver tissue is mechanically dissociated into small pieces and digested stirring in 10% FBS in RPMI containing 1mg/mL collagenase D and 0.06mg/mL DNAse I for 20 minutes. Hepatocytes are removed by centrifugation at 50*g* and CD34-enriched cells are obtained through density-gradient centrifugation with Lymphocyte Separation Medium (PromoCell) at 1000*g* without brake for 20 minutes. Finally, CD34^+^ HSPCs are isolated by positive selection using EasySep Human CD34 Positive Selection Kit II (Stemcell Technologies) following the manufacturer’s protocol. CD34^+^ HSPC isolation from cord blood is performed using EasySep Human Cord Blood CD34 Positive Selection Kit II (Stemcell Technologies) following the manufacturer’s protocol. Briefly, HSPCs are enriched by adding the RosetteSep cocktail to whole blood followed by centrifugation on a lymphoprep (Stemcell technologies) density gradient. Enriched mononuclear cells then undergo positive selection to purify CD34^+^ HSPCs. HSPCs are stored in 10% DMSO in FBS in liquid nitrogen.

### CRISPR editing

CD34^+^ human HSPCs are used immediately after isolation or recovered from frozen vials by culturing overnight in SFEMII medium (Stemcell Technologie) supplemented with 100ng/mL human SCF, 100ng/mL human TPO, 100ng/mL human FLT3L, 20ng/mL human IL-6 (cytokines from Peprotech), 0.75μM StemReginin 1 (Cayman Chemical), 500nM UM729 (Stemcell Technologies) at a density of 500,000 cells/mL. CRISPR ribonucleoprotein (RNP) is generated by combining 40pmol of Cas9 (IDT), 100pmol of sgRNA (Synthego) and PBS in 4μl for 10 minutes at room temperature. RNP is stable in 4°C. Since two or three guides are used per gene, a mix of sgRNA totaling 100pmol is used per reaction. HSPCs are concentrated by centrifugation and resuspended in 20μl P3 buffer (Lonza V4XP-3032), combined with CRISPR RNP and electroporated using the Lonza 4D Nucleofector using the DZ-100 program and p3 buffer. Cells are rested in warm medium for 30 minutes before engraftment in recipient MISTRG6 mice. For assessing editing efficacy by Inference of CRISPR Edit Analysis (40), DNA from edited cells are extracted using a H_2_O solution containing 50mM Tris, 1mM EDTA, 0.5% Tween-20 supplemented with 0.6mg/mL proteinase K incubated at 55°C for 3 hours or overnight and heat-inactivated at 95°C. Alternatively, DNA is extracted using the Quick-DNA microprep kit (Zymo Research). A region flanking the sgRNA cut sites is amplified by PCR and submitted for analysis on EditCo website (https://ice.editco.bio/). Guide sequences are as follows:

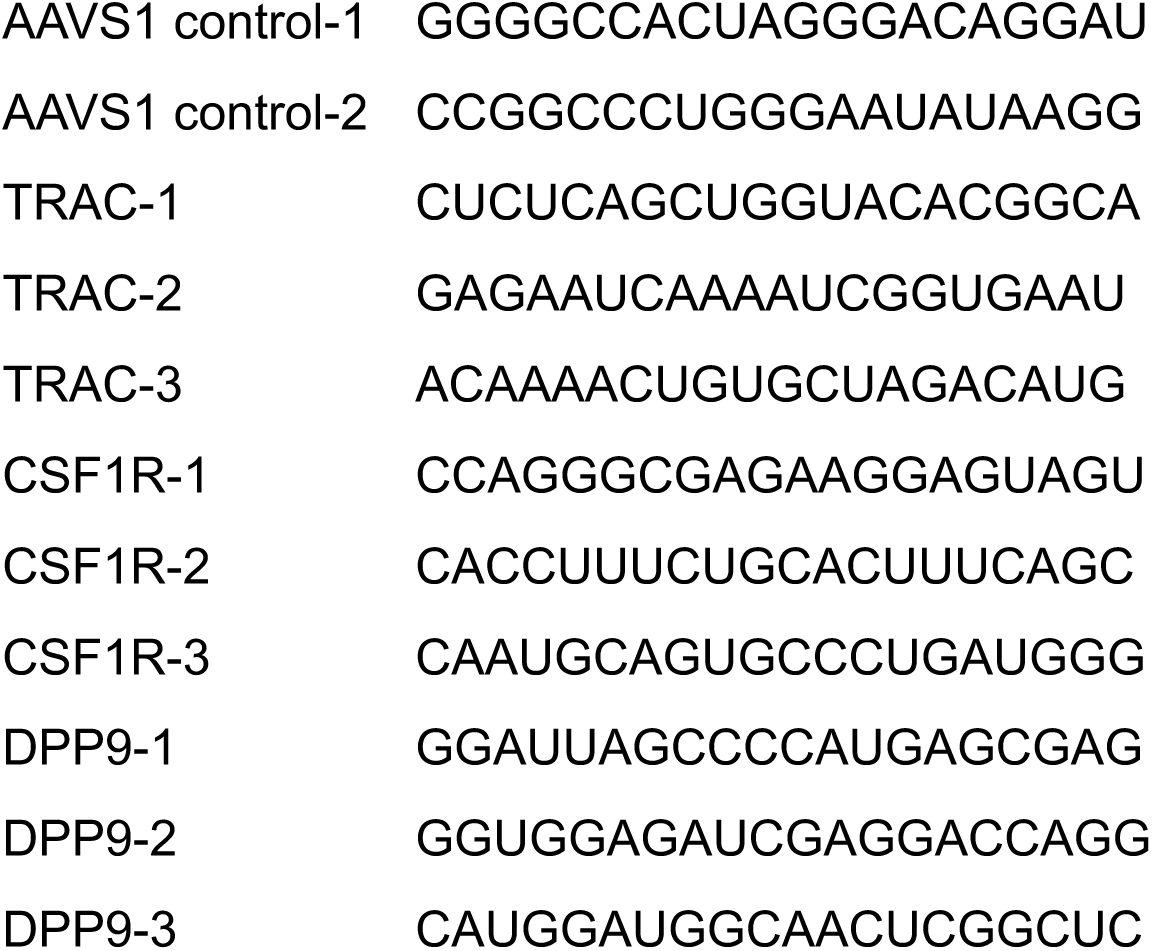
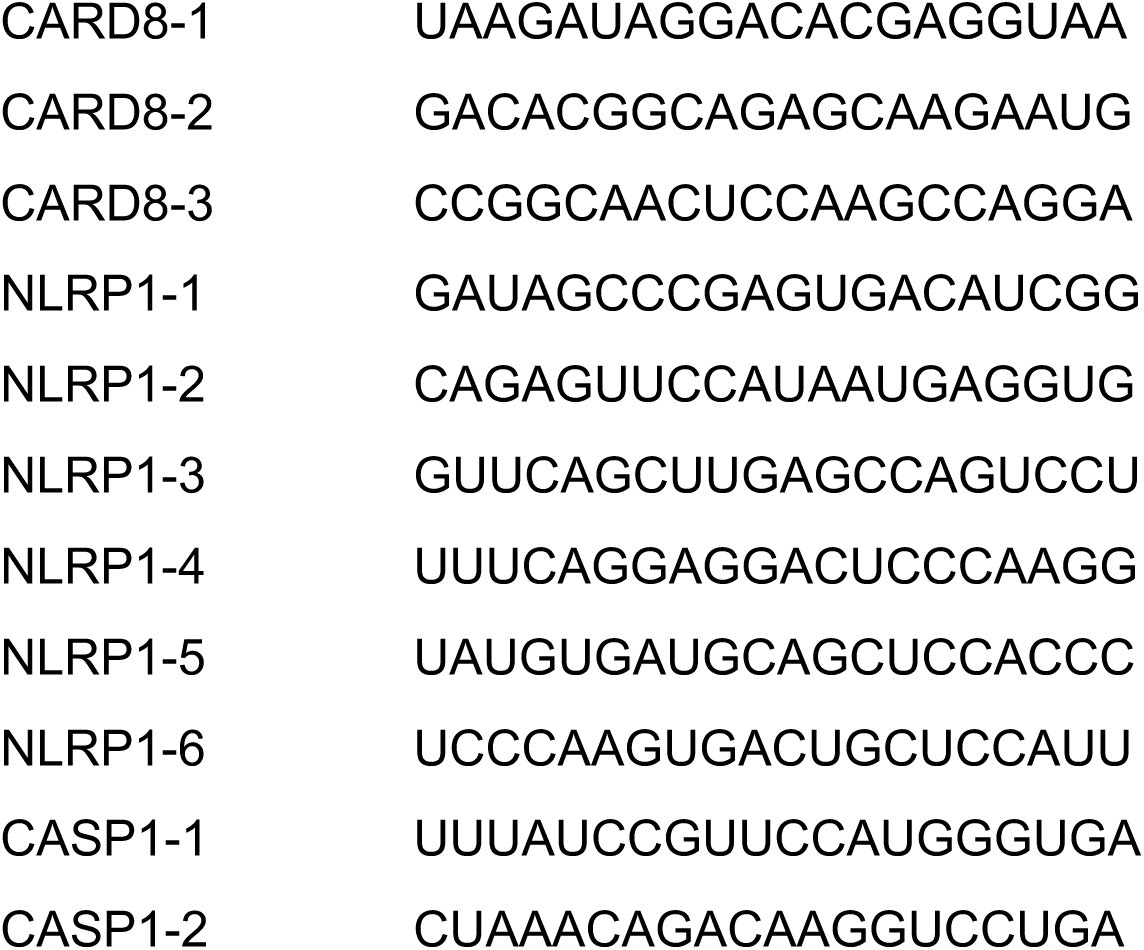

### iPSC cultures

human iPSC cell lines are derived in a previous study (75). Human iPSCs are grown as sparse aggregates in mTeSR plus media (Stemcell Technologies) and differentiated into hematopoietic progenitor cells (HPCs) using the STEMdiff hematopoietic kit (Stemcell Technologies) following the manufacturer’s protocol. Briefly, iPSC aggregates larger than 50μm are seeded at a density of 10-20 aggregates/cm^2^ in a 12-well plate. Aggregates are cultured in Base medium supplemented with medium A for 3 days, performing a half-medium change on day 2. Cells are cultured in Base medium supplemented with medium B from day 3 to day 12, performing a half-medium change on day 5 and on day 7, and on day 10. If needed, cells are stored in CS10 freezing media (Stemcell technologies) prior to in vivo engraftment.

### Cell isolation for flow cytometry and cell sorting

MISTRG6 mice are euthanized with 100% isoflurane. Blood was collected retro-orbitally into EDTA-containing solution for a final concentration of 4μm EDTA. Blood was directly stained with antibodies on ice. After washing, red blood cells lysis and fixation was achieved in one step using the RBC Lysis/Fixation Solution (Biolegend) following the manufacturer’s protocol. For cell isolation from the bone marrow, femurs are isolated using dissection tools and crushed using a mortar and pestle, then filtered through a 70μm filter. For cell isolation from the liver, liver is perfused. Immune cells are isolated using RPMI with 2% FBS and 0.25mg/mL collagenase II (Gibco), 0.1mg/mL DNAse I (grade ii, 10104159001 Roche) and 5mM calcium (Sigma). The same liver lobe is chopped into small pieces using scissors and digested stirring at 37°C for 30 minutes at 400rpm. Enzymes are quenched by adding 40mL of media, and hepatocytes are removed by centrifugation at 100*g* for 5 minutes twice. Immune cells from the supernatant are isolated through centrifugation at 500*g* for 5 minutes. Red blood cells are removed through ACK lysis. For cell isolation from the lung, the lung is digested with RPMI supplemented with 2% FBS, 1mg/mL collagenase D (Sigma) and 0.1mg/mL of grade II DNAse I (Roche). Surface antigen staining is as previously described(76). Briefly, single-cell suspensions are stained with live/dead stain in PBS on ice for 10 minutes, then stained for cell-surface antigens followed by washing in PBS supplemented with 2% FBS. When appropriate, cells are fixed with 2% paraformaldehyde in PBS until analysis. Cell sorting is performed on the BD FACSAria or Bigfoot Spectral Cell sorter. Flow cytometry analysis is performed using BD Symphony or Cytek Aurora.

The following antibodies were used: anti-human CD3 (OKT3), anti-human CD10 (HI10a), anti-human CD14 (HCD14), anti-human CD16 (3G8), anti-human CD19 (HIB19), anti-human CD20 (2H7), anti-human CD33 (WM53), anti-human CD34 (561), anti-human CD38(HIT2), anti-mouse CD45 (30-F11), anti-human CD45 (HI30), anti-human CD45RA (HI100), anti-human CD49f (GoH3), anti-human CD56 (HCD56), anti-human CD66b (G10F5), anti-human CD90 (5E10), anti-human CD123 (6H6), anti-human HLADR (L243). Live/Dead fixable viability dye yellow (Thermo Fisher Scientific) distinguished live and dead cells.

### Digital droplet PCR

DNA from cells is isolated using Quick-DNA microprep kit (Zymo Research). ddPCR is performed according to the Manufacturer’s protocol (Bio-Rad). Briefly, 25μl of ddPCR master mix is made with 2X ddPCR supermix for probes (no dUTP), 900nM primer, 250nM HEX or FAM probes. Droplets are generated from the qPCR master mix using the QX200 droplet generator (Bio-Rad). PCR is then performed on the C1000 Touch Thermal Cycler (Bio-Rad). Fluorescence of each droplet is read out using the QX200 Droplet Reader (Bio-Rad). To determine the fraction of knockout and control cells, a single DNA sample is fractionated into >20,000 droplets, and two PCR reactions are performed within each droplet. One PCR reaction with HEX fluorescent signal amplifies a reference *DPP9* sequence that is present in both control and *DPP9^-/-^* cell. The other PCR reaction (with FAM fluorescent signal) amplifies the *DPP9* knockout sequence targeted by the *DPP9* sgRNA such that control cells will have PCR amplification, but *DPP9^-/-^* cells do not. An absolute quantification of DNA copies for *DPP9* reference and knockout region is determined by quantifying the number of FAM^+^ or HEX^+^ droplets. Control cells have equal quantity of reference and knockout amplification, whereas *DPP9^-/-^* cells have only reference amplification. By comparing the ratio of reference and knockout amplification, we can quantify the presence of control and *DPP9^-/-^*cells.

### Colony forming unit assays and in vitro expansion assays

colony forming unit assay was performed as previously described (77). CD34^+^ cells were electroporated as above and seeded into a semi-solid MegaCult™-C (Stemcell Technologies, #04974) supplemented recombinant human cytokines: with 2.0 U/mL EPO, 10 ng/mL IL-3, 20 ng/mL IL-6, 50 ng/mL SCF, 50 ng/mL Thrombopoietin, 20 ng/mL G-CSF, 20 ng/mL M-CSF, and 10 ng/mL GM-CSF (ConnStem, Inc., USA). 13 days later, colonies were labelled by adding 4 tests per plate of antibodies targeting CD14 (M5E2), CD41a (HIP8), CD66b (6/40c), CD235a (HI264) (Biolegend 301830, 303704, 392904, 349114) diluted in 300uL PBS to detect colonies with monocyte, megakaryocyte, granulocyte, and erythrocyte lineages, respectively. Wells were imaged the next day with a Molecular Devices Image Xpress Micro 4 (IXM) high content microscope to produce high-resolution whole-well scans at 10X magnification. For colony forming unit assay of sorted HSCs, single CD90+ CD45RA-CD34+ CD38-CD45+ HSCs were sorted into 384 well plates containing a custom liquid format of the semi-solid media used above by replacing the collagen supplement with 0.12N HCl in dH2O. After 14 days, each well was stained by adding the same antibody cocktail as for the semi-solid format diluted in 5µL of PBS per well 24 hours prior to imaging or flow cytometry readout. For clonal expansion assay, single cells of HSCs were sorted into 384 wells containing SFEMII medium (Stemcell Technologie) supplemented with 100ng/mL human SCF, 100ng/mL human TPO, 100ng/mL human FLT3L, 20ng/mL human IL-6 (cytokines from Peprotech), 0.75μM StemReginin 1 (Cayman Chemical), 500nM UM729 (Stemcell Technologies) for 7 days. At each experimental endpoint, cells were acquired on a BD LSRFortessa flow cytometer.

### Single-cell RNA sequencing

Lin^-^ CD34^+^ human cells were sorted from engrafted bone marrow. Cells were hashed with 0.5μg of Totalseq anti-human hashing antibody (Biolegend B0251 for control and B0252 for *DPP9^-/-^*). Approximately 10,000 hashed cells were encapsulated into droplets using the 10x Chromium GEM platform. Single-cell RNA-seq libraries were generated using the Chromium Next GEM Single Cell 3′ Reagent Kits v3.1 (10x Genomics) according to the manufacturer’s instructions and sequenced on an Illumina NovaSeq system. Raw sequencing data were processed using Cell Ranger (v7.1.0) and aligned to the GRCh38-3.0.0 reference genome. The resulting filtered gene–barcode matrices were imported into R (v4.2.3) and analyzed using Seurat (v5.0.1). Wild-type (WT) and knockout (KO) cells were demultiplexed based on hashing antibody signals using the HTODemux() function. Cells were then filtered to exclude those with mitochondrial gene expression greater than 25%, followed by data normalization, scaling using ScaleData(), and principal component analysis (PCA). Datasets were subsequently merged, clustered, and visualized using Uniform Manifold Approximation and Projection (UMAP). Differential expression was performed using Seurat’s FindMarkers function. Pathway enrichment analysis was performed using Ingenuity Pathway Analysis (Qiagen).

### Statistical analyses

Data are routinely shown as mean ± SD. Statistical significance was determined by two-tailed Student’s t-test using GraphPad Prism 9.0. *P < 0.05, **P < 0.01, ***P < 0.001, and ****P < 0.0001 unless stated otherwise. In some figures, relative cell numbers are used instead of absolute numbers to normalize the cell number variations between different tissue donors. Relative cell numbers is given as the fold change of each individual animal relative to the mean of the control group within each experimental cohort.

### Study Approval

All procedures were carried out following protocols approved by the Yale Institutional Animal Care and Use Committee. This study is not considered Human Subjects Research.

## Data availability

Values for all data points in graphs are included in the Supporting Data Values file. Single-cell RNA-seq data generated in this study (GSE332722) have been deposited in Gene Expression Omnibus. All data needed to evaluate the conclusions in the paper are present in the main text or supplementary materials.

## Author contributions

The project was conceptualized by TX, JRB, AH. TX, JRB, MC, AH, YT, CL, FZ, MC, HNB performed experiments. QW, LS, ES provided key reagents. KB, DSK, RAF provided funding. RAF provided supervision of entire project. TX wrote the original manuscript with substantial input from AHN, which was edited by all authors.

## Funding

This work is the result of NIH funding, in whole or in part, and is subject to the NIH Public Access Policy. Through acceptance of this federal funding, the NIH has been given a right to make the work publicly available in PubMed Central.

NIH R21AI178249 and Howard Hughes Medical Institute to Richard A Flavell

American Cancer Society PF-25-1435117-01-PFCBI to Tianli Xiao

NIDDK U54DK106857 to Diane S Krause

## Acknowledgement

We thank P.Clarke for her help with ddPCR; J. Alderman for logistical support; C. Hughes for animal logistical support. This research was supported by the Yale University Reproductive Sciences Biobank HIC#12696, a component of the Department of Obstetrics, Gynecology & Reproductive Sciences, Yale School of Medicine, New Haven, CT. We thank Yale Flow Cytometry for their assistance with cell sorting and flow cytometry analysis. The Core is supported in part by an NCI Cancer Center Support Grant # NIH P30 CA016359. The BD Symphony was funded by shared instrument grant # NIH S10 OD026996.

## Figure legends

**Supp Fig. 1.**
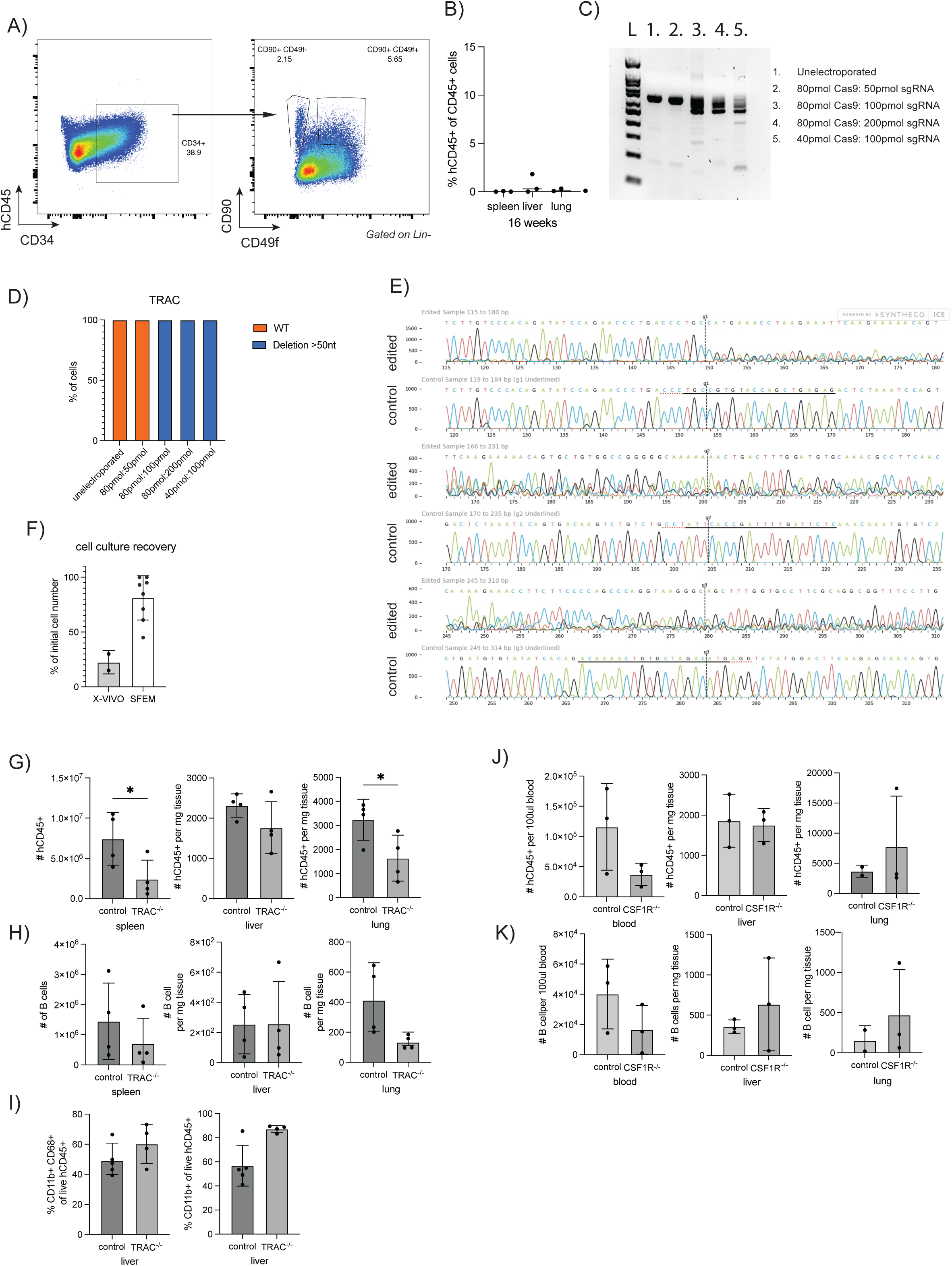
(**A**) Markers of iPSCs following hematopoietic stem cell differentiation following manufacturer’s protocol (Stemcell technologies). (**B**) Neonatal engraftment of iPSC-derived hematopoietic stem cells in MISTRG6 mice. hCD45^+^, human CD45^+^. (**C**) primary CD34+ human HSPCs are electroporated with varying quantity of Cas9 and sgRNA against the TRAC locus. PCR amplicon around the edited region. L: Ladder. (**D**) Inferred Editing Efficacy (Synthego) based on Sanger traces from samples in panel C). (**E**) Sanger traces around the *TRAC* locus around sgRNA 1 cut site (g1), sgRNA 2 cut site (g2), sgRNA 3 cut site (g3). (**F**) CD34^+^ human HSPCs are cultured overnight in X-VIVO 15 or SFEM media and the number of cells recovered the next day is quantified. (**G**) number of human CD45+ cells, (**H**) human B cells, and (**I**) frequency of human myeloid cells 12-16 weeks after engraftment with control or *TRAC^-/-^* CD34+ HSPCs in MISTRG6 mice. (**J**) number of human CD45+ cells and (**K**) human B cells 9 weeks after engraftment with control or *CSF1R^-/-^* CD34+ HSPCs in MISTRG6 mice.

**Supp Fig. 2.**
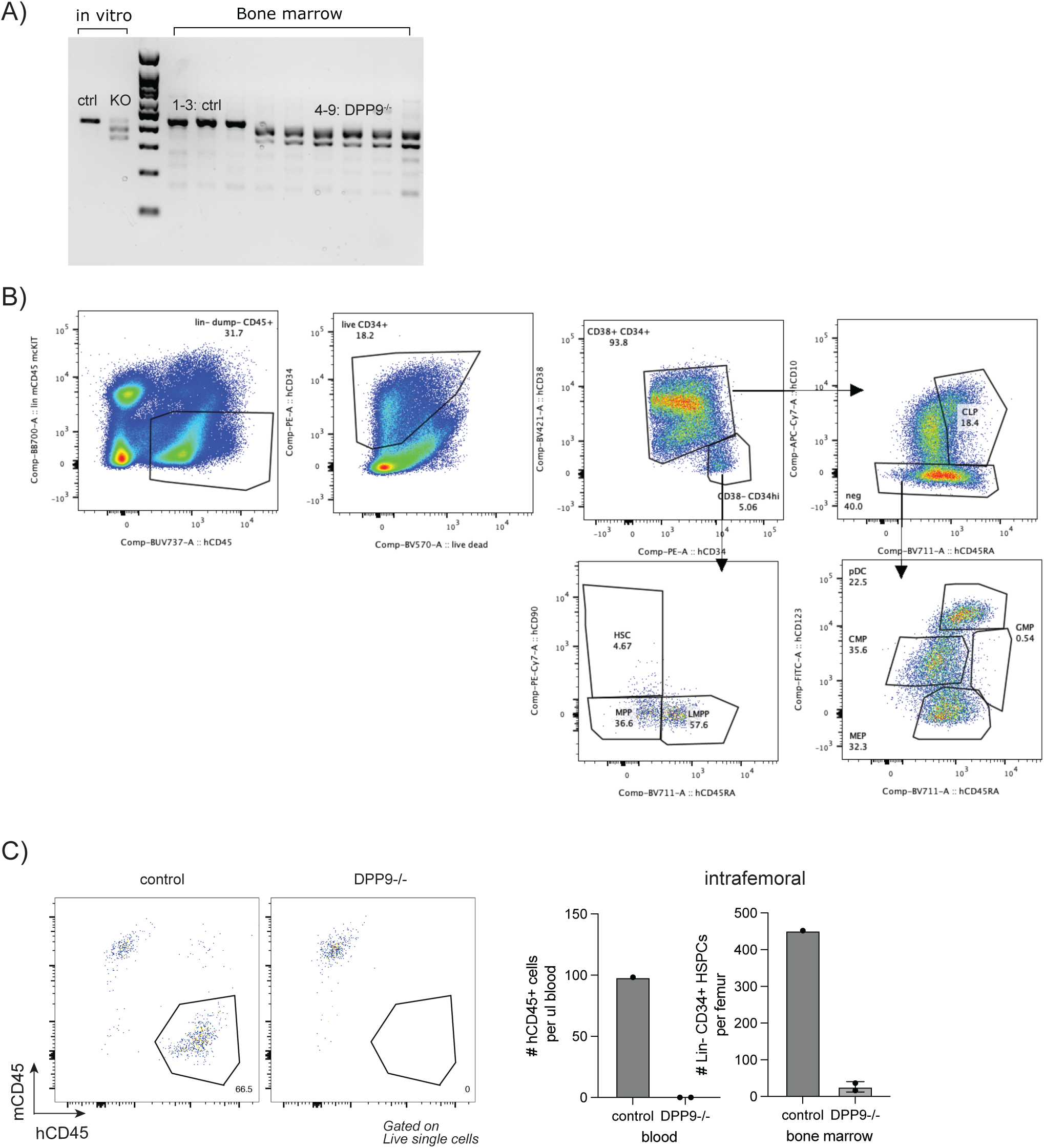
(**A**) PCR amplification of the *DPP9* sgRNA targeting locus. Edited cells have a smaller amplified band due to truncation deletion caused by CRISPR editing. CD34+ cells are either engrafted in vivo or kept in culture in vitro culture for 3 days. DNA from bone marrow 9 weeks after engraftment. (**B**) Gating strategy for bone marrow hematopoietic stem and progenitor populations. (**C**) Control or *DPP9^-/-^* cells are injected intrafemorally for 16 weeks, cell numbers are assessed in the blood and bone marrow.

**Supp Fig. 3.**
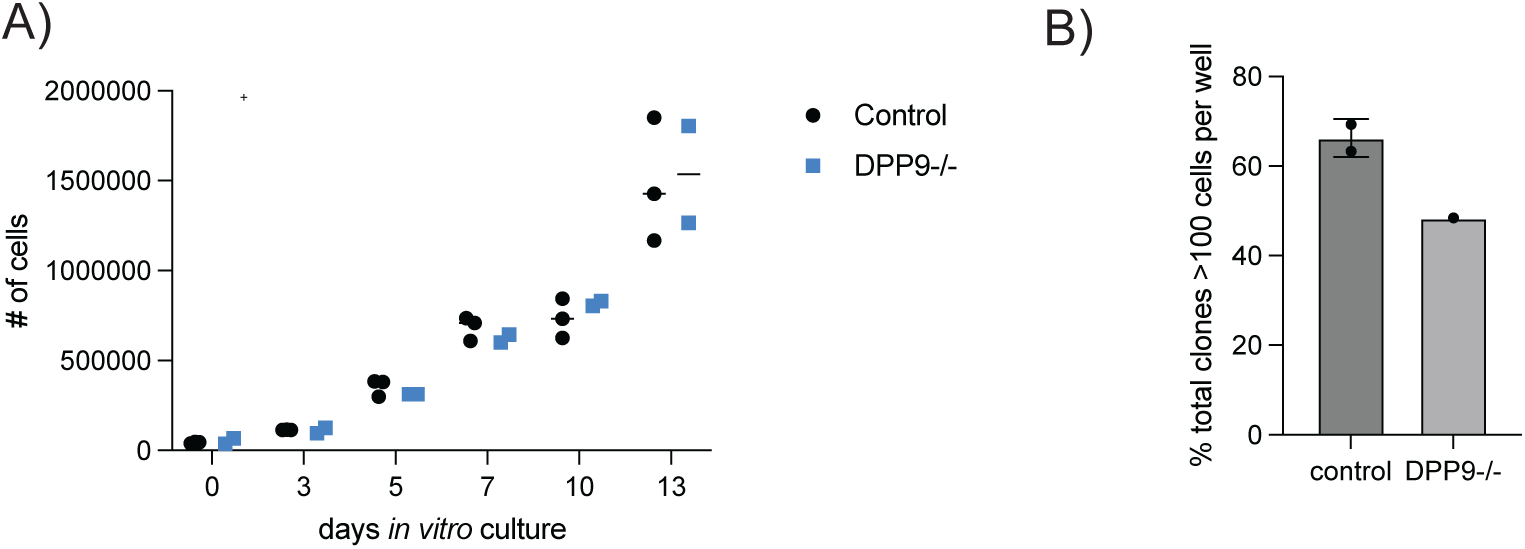
(**A**) In vitro expansion culture of control and *DPP9^-/-^* CD34^+^ HSPCs. (**B**) Frequency of clones that expanded to more than 100 cells at the end of a 7-day expansion of HSCs following experimental scheme in Figure 3D.

**Supp Fig. 4.**
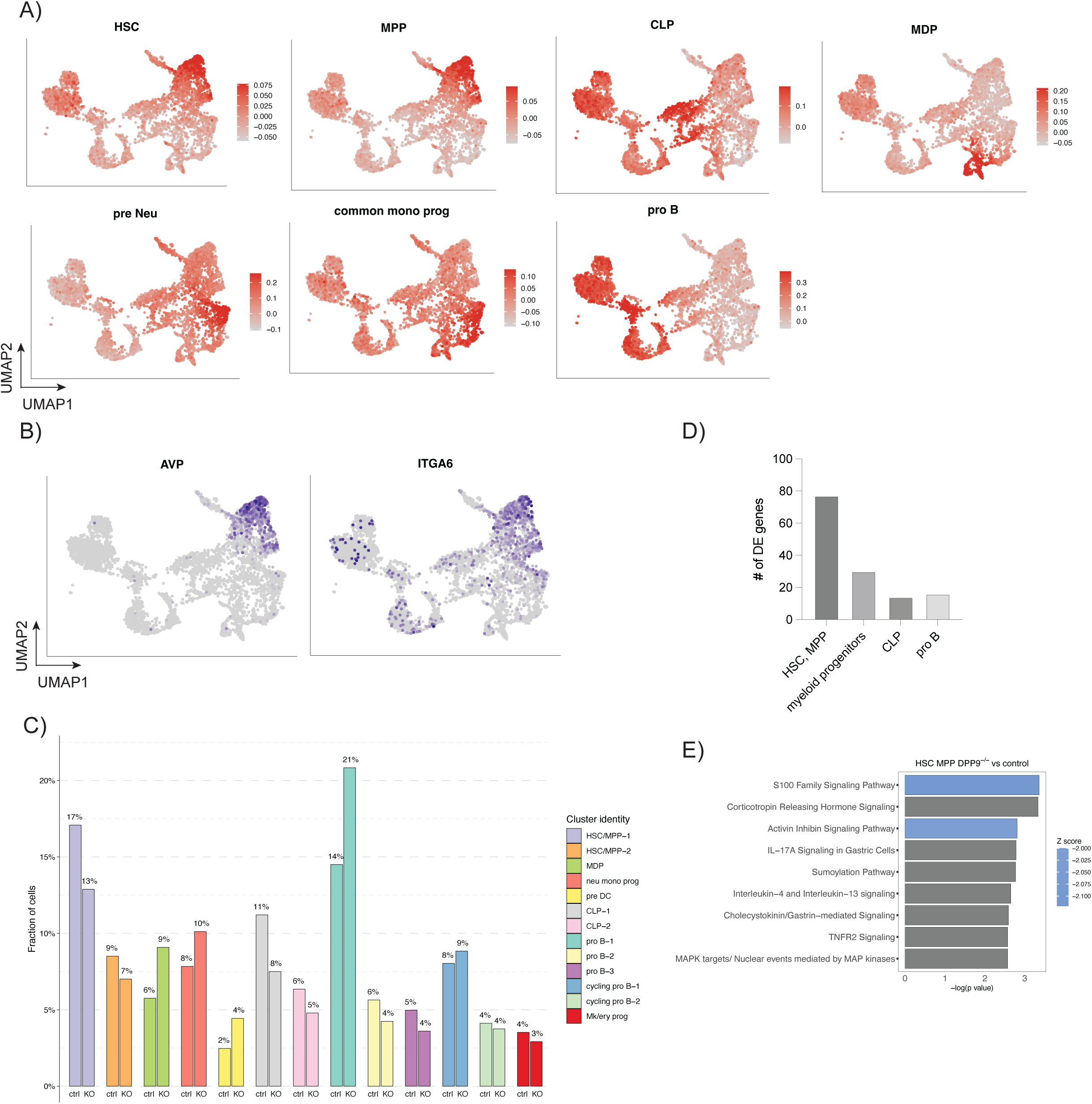
(**A**) Overlay of various gene signatures of different human bone marrow populations (49) onto scRNA-seq dataset. (**B**) Expression of *AVP* and *ITGA6* encoding CD49f marking HSC populations. (**C**) Frequency of cells within each cluster for control or *DPP9^-/-^* cells. WT, control. KO: *DPP9^-/-^*. (**D**) number of differentially expressed (DE) genes defined as FDR <0.1 for each cluster. HSC, MPP includes HSC/MPP-1 and HSC/MPP-2, myeloid progenitors include MDP, neu mono prog, pre DC. CLP includes CLP-1, CLP-2. Pro B includes pro B-1, pro B-2, pro B-3. (**E**)Ingenuity Pathway Analysis of differentially expressed genes found between *DPP9^-/-^*and control HSC/MPP clusters, DE genes defined as FDR <0.1.

**Supp Fig. 5.**
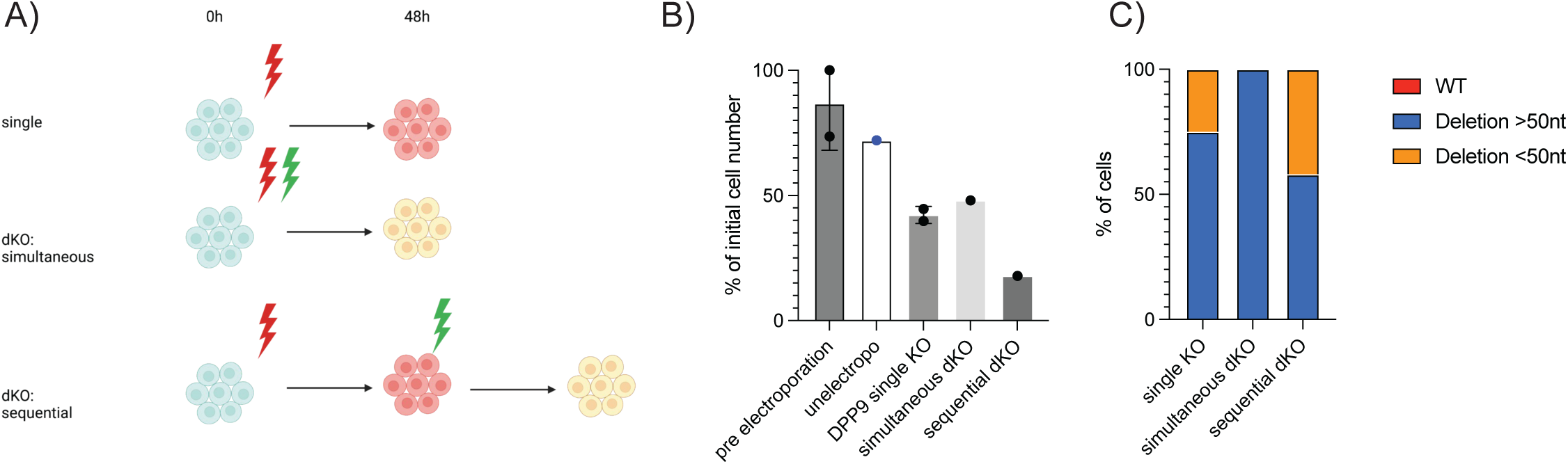
(**A**) Schematic for sequential or simultaneous double KO. (**B**) Frequency of initial cell number after electroporation. (**C**) gene knockout efficiency of *DPP9* by Sanger sequencing.

